# Outcome-Driven Microscopy: Closed-Loop Optogenetic Control of Cell Biology

**DOI:** 10.1101/2024.12.12.628240

**Authors:** Josiah B. Passmore, Alfredo Rates, Jakob Schröder, Menno T. P. van Laarhoven, Vincent J. W. Hellebrekers, Henrik G. van Hoef, Antonius J. M. Geurts, Wendy van Straaten, Wilco Nijenhuis, Florian Berger, Carlas S. Smith, Ihor Smal, Lukas C. Kapitein

## Abstract

Smart microscopy is transforming biological imaging by integrating real-time analysis with adaptive acquisition to enhance imaging efficiency. Whereas many emerging implementations are event-driven and focus on on-demand data acquisition to reduce phototoxicity, we here present ‘outcome-driven’ microscopy, which combines smart microscopy with optogenetics to achieve subcellular spatiotemporal control of biology to predefined outcomes. We validate this approach using light-based control of cell migration and nucleocytoplasmic transport, and demonstrate unprecedented spatiotemporal control over cellular behaviour.

## MAIN

Biological imaging is constantly evolving, with ‘smart’ microscopes paving the way for efficient, adaptive workflows. A smart microscopy platform performs real-time analysis, adjusting acquisition parameters on the fly^1–6^. Typical implementations of such real-time feedback loops use the information from the acquisition to change microscope behaviour, such as imaging modalities, speed, or dimensions. This allows the microscope to adapt to the sample and minimise imaging that does not contribute to the biological question, thus preserving sample health by reducing phototoxicity and improving efficiency^1,7,8,5^. As such, the majority of smart microscopy implementations record the sample passively, for example to track objects^9–13^, continually optimise imaging parameters^10,14–17^ or, in the case of event-driven microscopy, adjust modalities in response to specific events^18–20^.

Beyond observation of a biological process, microscopy can also facilitate active control. Most notably, optogenetics provides a platform for inducible control of biological systems across scales^21–25^ using light, with a dose-response relationship^26–29^ and a rapid localised response of cells to spatial light patterns^30–33^. Optogenetics has been previously combined with closed-loop smart microscopy platforms in a passive manner, with the controller adjusting hardware on the fly through automated segmentation or tracking^34–38^. However, the control of a biological system available through optogenetics provides the opportunity to extend feedback control to optimise not only hardware in passive observation, but also actively control biological processes themselves to achieve user-defined outcomes^39,22^. Indeed, combinations of closed-loop control and optogenetics have previously enabled such ‘outcome-driven’ research, adjusting light intensity or patterns on-the-fly *in vivo*^40–47^, or in cultured cells^48–63^ as demonstrated in cybergenetics^64^ – most often with specific custom platforms designed for intensity optimisation at the population or whole-cell level for gene expression. As such, none of these examples fully harness the potential of optogenetics to achieve automated subcellular control of cellular dynamics. Such control over biology has been a long-term goal of cell biological research^65^.

Here, we present a generalised platform for outcome-driven microscopy that uses optogenetics and real-time feedback to achieve automated spatiotemporal control of subcellular cell biology. This platform can control both the spatial patterning and intensity of light and adjusts them on-the-fly to bring biological systems to predefined outcomes, as demonstrated by long-term guidance of cell migration over predefined paths and by the controlled titration of protein levels in the cytosol or nucleus.

First, we built a modular smart microscopy platform, with interchangeable modules for microscope communication strategies, image processing pipelines, and control approaches (Extended Data Fig. 1 and Supplementary Text 1). This platform can be adapted for a variety of experiments by adjusting the modules to fit specific cases. For our implementation of outcome-driven microscopy, a feedback loop is established whereby the microscope captures an image for processing, and based on the information extracted from the image, the control module optimises specific biological processes to achieve predefined outcomes by adjusting the illumination parameters of a spatial light modulator.

To test the capabilities of outcome-driven microscopy, we used cell migration as our first proof-of-concept. Control of cell polarisation and migration using optogenetics is well established through the use of photoactivatable Rho GTPases or recruitment of upstream effectors to the plasma membrane^66–72^ (Fig. 1a). Light directed migration has been achieved by gradient illumination^70^, manually updating the area of illumination (AOI)^67–69,73,74^, or with smart microscopy, using live segmentation with tracking to achieve constant illumination in the same cellular area^34–36^. We reasoned that outcome-driven microscopy could push this further, and constantly update the AOI to direct cells to predefined migration paths. This approach would allow us to overcome stochastic variations such as differences in protein expression levels, morphology, or response times, with the controller ensuring that cells stay on the predefined track. To establish this, we used feedback on the position of the cell centroid relative to a path made up of setpoints, combined with a trajectory tracking controller to selectively illuminate the cell in the region closest to the next selected setpoint (Fig. 1b and Supplementary Text 1). For reliable segmentation, we used the pre-trained AI segmentation method Segment Anything Model (SAM)^75^, with a custom extension of its interface for tracking (Supplementary Text 2). Using the migratory cell line HT1080 stably expressing an optogenetic construct to recruit the RAC1 effector TIAM1 to the plasma membrane^72^, we found that we could reliably guide cells around a specific path multiple times for >10 hours (Fig. 1c and Supplementary Video 1), maintaining a relatively consistent speed (Fig. 1d and Extended Data Fig. 2a). Quantifying the distance between the centroid and the desired path (path deviation) revealed that the centroid of the cell was kept within 2.5 µm of the path (Extended Data Fig. 2b), demonstrating precise control of directed cell migration.

**Fig. 1:**
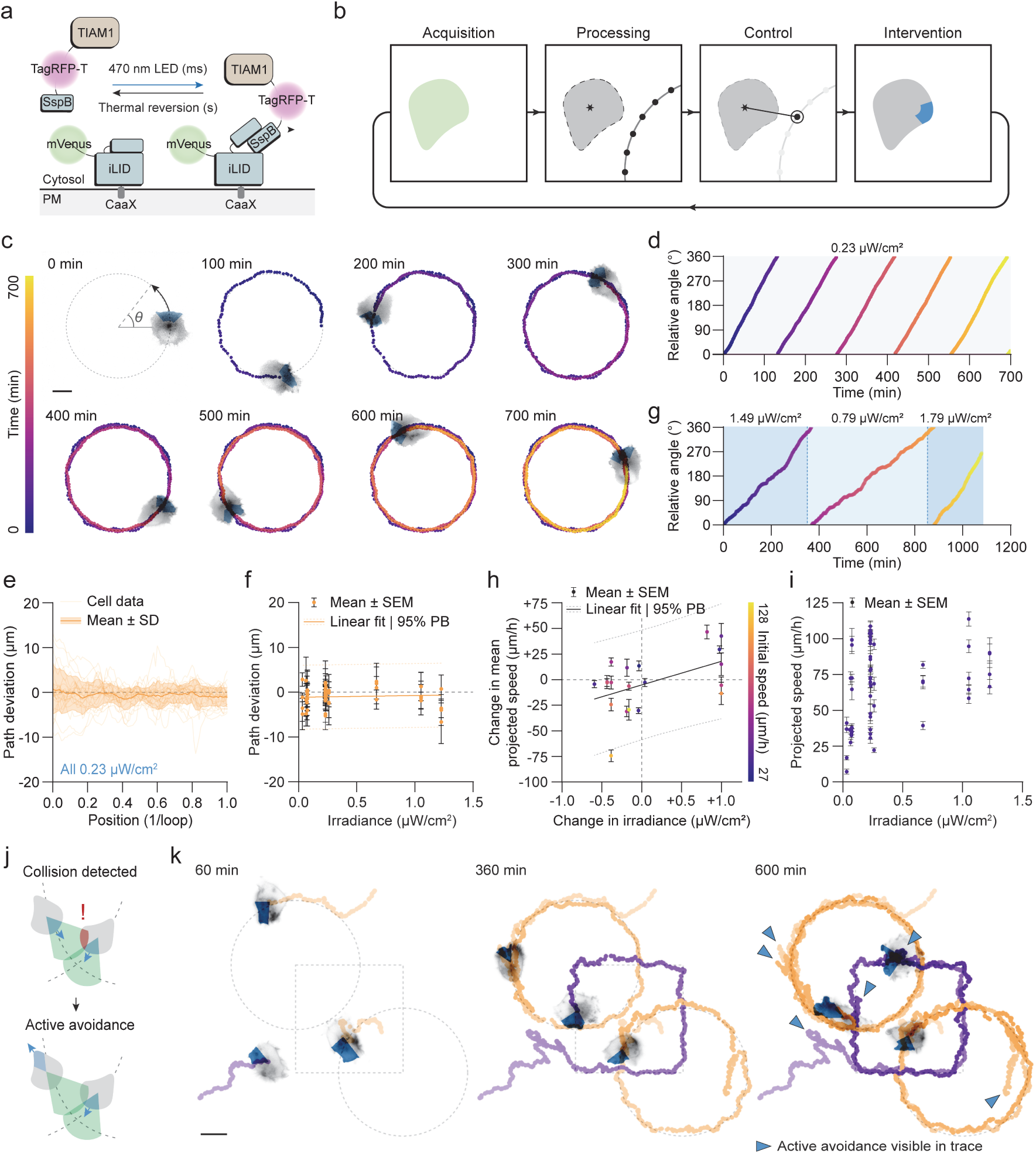
Outcome-driven microscopy enables directed cell migration. **a**) Schematic of the construct used for optogenetic recruitment of TIAM to the plasma membrane^71^. **b**) Outcome-driven microscopy approach to control of directed cell migration using a direct orientation-correction controller. **c**) A HT1080-TIAM cell (grey) controlled using outcome-driven microscopy to illuminate a specific region of the cell (blue) and induce migration in a circle path (dashed line). The cell centroid was tracked and marked (colour bar). Scale bar: 10 µm. **d**) The angle of the cell centroid in **c** over time, relative to the starting position. **e**) The path deviation of all cells guided to circle paths, with a constant irradiance of 0.23 µW/cm^2^, separated by individual loops. n = 24 cells. **f**) Average path deviation of all cells guided to circle paths with various irradiance levels. PB = prediction band. n = 58 cells. **g**) The angle of the cell centroid relative to the starting position for a cell with changing irradiance during the acquisition. **h**) The change in mean projected speed for cells that were subjected to a change in irradiance between individual loops during guidance. Initial speed denotes the mean projected speed of the cell in the previous loop. PB = prediction band. n = 20 cells. **i**) Average projected speed of each individual loop of controlled cells, per irradiance. n = 58 cells. **j**) Schematic of the active avoidance system for multi-cell outcome-driven control of directed cell migration. **k**) Three cells were simultaneously controlled to individual overlapping paths, with an active avoidance system as in **j** pulling back cells to ensuring collisions were avoided. Cell centroids were tracked and overlaid (orange and purple), with blue arrowheads showing points in the track where successful avoidance is visible. Scale bar: 20 µm.

Outcome-driven microscopy provides the opportunity to control multiple cells in the same manner to the same setpoint, while also considering cell-cell variation. Indeed, when controlling multiple cells with the same constant LED irradiance (0.23 µW/cm^2^), we observed some variation in cell speed, independent of the expression level of TIAM1 (Extended Data Fig. 2c-d). Despite this heterogeneity, the controller was able to keep all cells within 5 µm of the desired path throughout the experiment (Fig. 1e). Path deviation was consistently low between cells, even at the lowest irradiance (Fig. 1f and Extended Data Fig. 2e), indicating that even a small degree of optogenetic activation is sufficient to maintain a consistent signalling pathway and guide cells to specific paths with outcome-driven microscopy.

Next, we asked if, despite cell-cell variation in speed when keeping irradiance constant, we could modulate cell speed by adjusting irradiance over time, due to the dose-response relationship of optogenetics. To evaluate this relationship, we guided cells around a circular path, and adjusted irradiance in a stepwise manner, changing the value for each complete or half loop (Fig. 1g and Supplementary Video 2). Indeed, a greater variation in speed is seen when guiding cells with different light levels (Extended Data Fig. 2f), and notably, we observed some degree of irradiance-dependence of relative cell speed (Fig. 1g, h), demonstrating that adjusting irradiance can speed up or slow down cells. Interestingly, the inherent variability in the baseline speed of unstimulated, undirected cells (Extended Data Fig. 2g) prevented us from consistently achieving specific absolute speed setpoints at fixed irradiance levels (Fig. 1i). We observed that cells that are initially faster are more capable of decreasing their speed, and cells that initially move slow are easier to speed up. This indicates a saturation point, suggesting that speed can only be modulated within a certain global range, likely due to inherent physiological constraints or saturation of the signalling pathway.

To demonstrate the high level of control that we can achieve and to push the limits of outcome-driven control of cell migration, we adapted our platform for outcome-driven microscopy to control multiple cells simultaneously, each on a different path. Additionally, to avoid collisions we added coordination between cells by incorporating a lookahead, whereby the cells will be pulled back in the direction opposing their initial movement if an imminent collision is detected (Fig. 1j). We observed that we could effectively guide multiple cells to their respective paths with minimal error, and also effectively stall them to avoid collisions (Fig. 1k and Supplementary Video 3).

In addition to outcome-driven microscopy aimed at controlling whole-cell dynamics with spatial illumination patterns, we next set out to establish a subcellular proof-of-concept and explore an intensity-driven relationship in depth to further demonstrate flexibility in control strategy. The blue light-induced nuclear export system (LEXY)^76^ has a rapid light-response, and induces striking subcellular localisation differences with a dose-response relationship (Fig. 2a). We hypothesised that we could achieve full control of protein levels within the cytosol and nucleus using outcome-driven microscopy by adjusting illumination irradiance on the fly, placing irradiance under control of a proportional-integral-derivative (PID) controller^77^ (Fig. 2b). The controller is optimised to minimise the error of the intensity in either the nucleus or cytosol in comparison to a setpoint. To reliably segment both the nucleus and the cytosol, we used our custom interface for the pre-trained SAM^75^ and implemented a temporal filter to resolve timepoints with low contrast between the intensity of the nucleus and the cytosol (Supplementary Text 2).

**Fig. 2:**
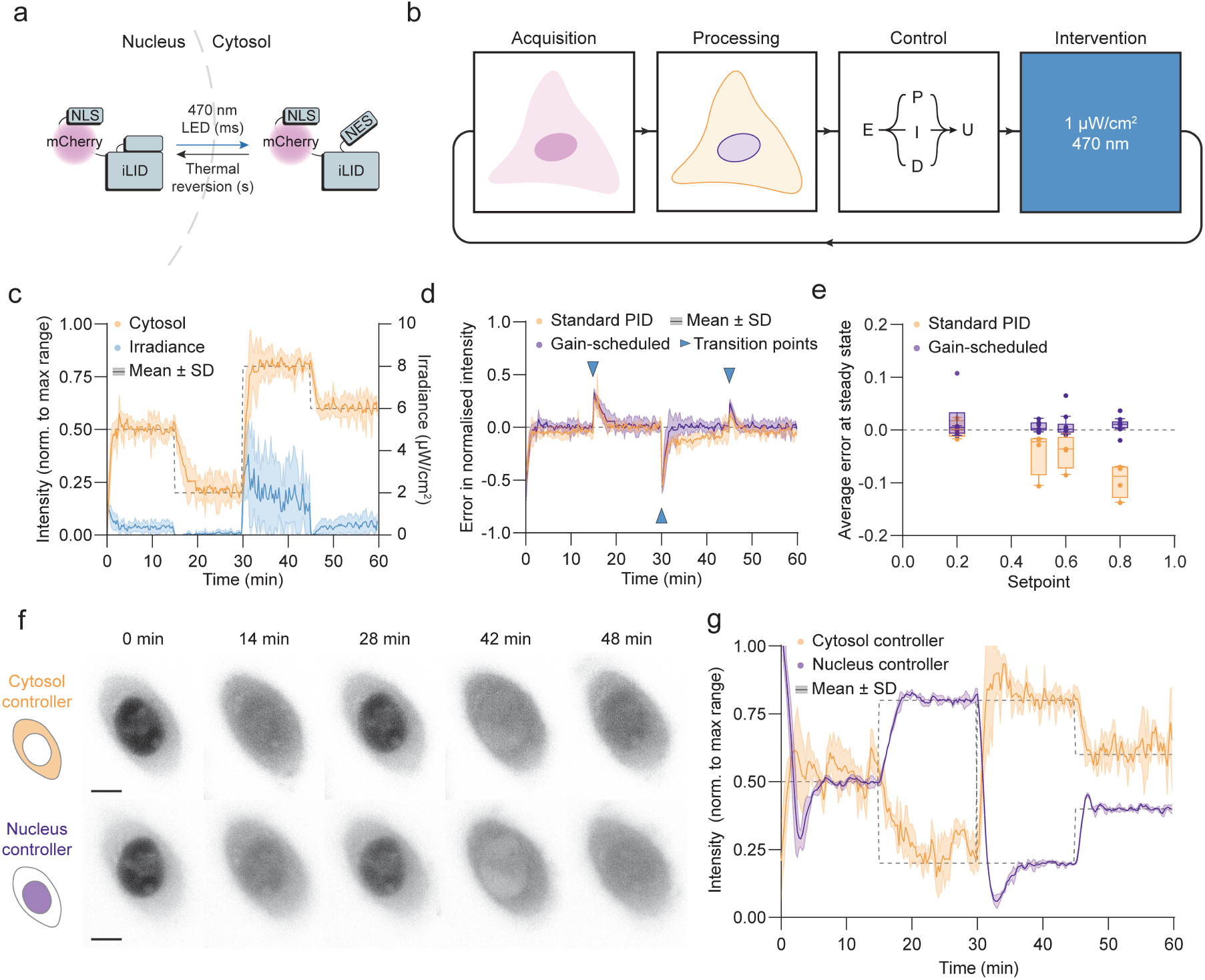
Outcome-driven microscopy enables well-controlled titration of protein levels in the nucleus or cytosol. **a**) Schematic of the LEXY construct^76^ used for optogenetic uncaging of a nuclear export sequence. **b**) An outcome-driven microscopy approach to control of nucleocytoplasmic transport using a proportional-integral-derivative (PID) controller. **c**) U2OS-LEXY cells were controlled using outcome-driven microscopy to bring cytosolic intensities (orange) to specific setpoints (dashed line) by modulating irradiance over time (blue) using a gain-scheduled PID controller. Intensities are represented normalised to the maximum dynamic range. n = 10 cells. **d**) The error in normalised intensity (intensity–setpoint) for cells guided to the same setpoints as in **c**, with the standard and gain-scheduled PID controllers. Transition points (changes of setpoint) are shown with blue arrowheads. N = 4 cells (standard), n = 10 cells (gain-scheduled). **e**) The distributions of average error at steady state for each setpoint, for both standard and gain-scheduled PID controllers. n = 4 cells (standard), 10 cells (gain-scheduled). **f-g**) a U2OS-LEXY cell (grey) was controlled using controllers for cytosol intensity (orange) and nucleus intensity (purple) to various setpoints (dashed lines). n = 3 runs for each controller. Examples in **f** show the first run of each controller. Scale bars: 10 µm.

First, we measured cell-cell variation in nucleocytoplasmic transport rates to inform our control strategy. In U2OS cells stably expressing LEXY, we observed that the expression level was a major contributor to cell-cell variation, with higher expressing cells having a lower rate of export (Extended Data Fig. 3a). Interestingly, we find that import rates do not correlate with the expression level, as well as the steady-state ratio of intensities between the two compartments (Extended Data Fig. 3b-c). After tuning a basic PID controller based on a model of this system (Supplementary Text 3), we were able to control cytosolic intensity to desired setpoints, and the error between measured intensity and setpoint was consistently <10% (Extended Data Fig. 3d-f). However, we found that due to our design constraints aimed at minimising overshoot, these model-derived gains were not universal enough to deal with the nonlinear dynamics and cell-cell variation, especially when dynamically changing setpoints across the whole range of cell intensities (the operating range). In these cases, we often observed suboptimal behaviour such as steady-state offsets, depending on the cell (Extended Data Fig. 3g).

Therefore, we improved out controller by including gain scheduling, which adjusts PID gains as a function of a scheduling variable^78^ – in our case, measured intensity – allowing gains to be optimised across the whole operating range. We implemented a gain-scheduled PID controller by first selecting a set of gains for various operating points, using our first-principle model of the system (Supplementary Text 3). We then fine-tuned the controller to find an optimal value-gain matrix. Using this gain-scheduled controller, we were able to control cells for various setpoints within the whole operating range to a high degree of accuracy, overcoming cell-cell variation and stochasticity, and outperforming the standard PID controller across the entire range (Fig. 2c-e).

To demonstrate flexibility in control strategies, and because we envision applications of outcome-driven control of nucleocytoplasmic transport that would benefit from control of nuclear intensity, we also developed a controller to modulate intensity levels in the nucleus (Supplementary Text 3). To compare the two controllers and to show robustness of control, we ran both the cytosol and nucleus controllers in the same cell, repeating three times for each controller (Fig. 2f-g and Supplementary Video 4). Both controllers were highly reproducible, and were able to bring intensities to the predefined setpoints with remarkably similar irradiance levels, and with minimal error (Extended Data Fig. 3h-i). Interestingly, we see that the nucleus controller is less noisy than the cytosol controller, likely because the average nucleus intensity is more representative of the real concentration than the average cytosol intensity is for the cytosol, due to the dynamic three-dimensional shape of the cell, the distribution of the marker across the cytosol, and changes of the segmentation through time. These experiments pave the way for the controlled titration of protein levels in nucleus or cytosol, which could for example be used to explore the dose-response functions of different transcription factors.

In summary, we have established a proof-of-concept for the robust integration of smart microscopy and optogenetics to overcome cell-cell variation, bring cells to predefined outcomes, and achieve precise, outcome-driven control of microscopy experiments. This approach is also easily applicable to other existing microscope communication platforms that are used for smart microscopy^10,79–81^. Advancing the field of smart microscopy by integrating active control of cell biology through optogenetics, we provide a platform for reproducible, cell-cell variation-independent, interrogation of biological processes. This outcome-driven approach can be further expanded into multicellular control and/or 3D environments, in order to provide insight into complex biological interactions and control morphodynamical transitions.

## EXTENDED DATA

**Extended Data Fig. 1:**
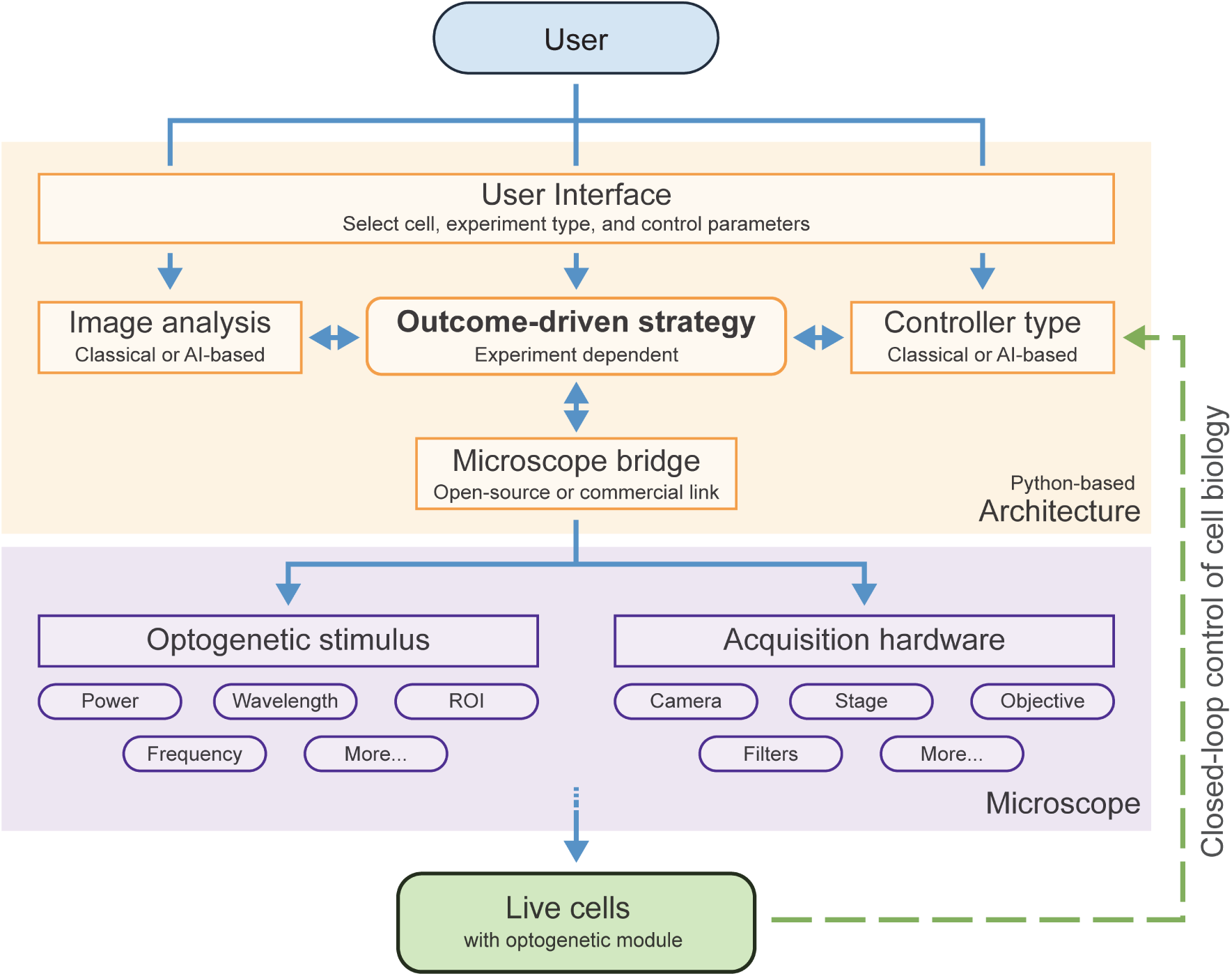
A modular platform for smart, outcome-driven microscopy. A python-based architecture (orange) consisting of user-adjustable modules for microscope bridging, image analysis and feedback control, coordinated by a user-defined outcome-driven strategy. Microscope hardware (purple) is controlled by the architecture bridge, to adjust optogenetic stimulus through a spatial light modulator, change acquisition parameters, and acquire an image. The acquired image of the modified sample is fed back into the architecture at each timepoint to ultimately bring cells to a specific outcome. For a complete description, see Supplementary Text 1.

**Extended Data Fig. 2:**
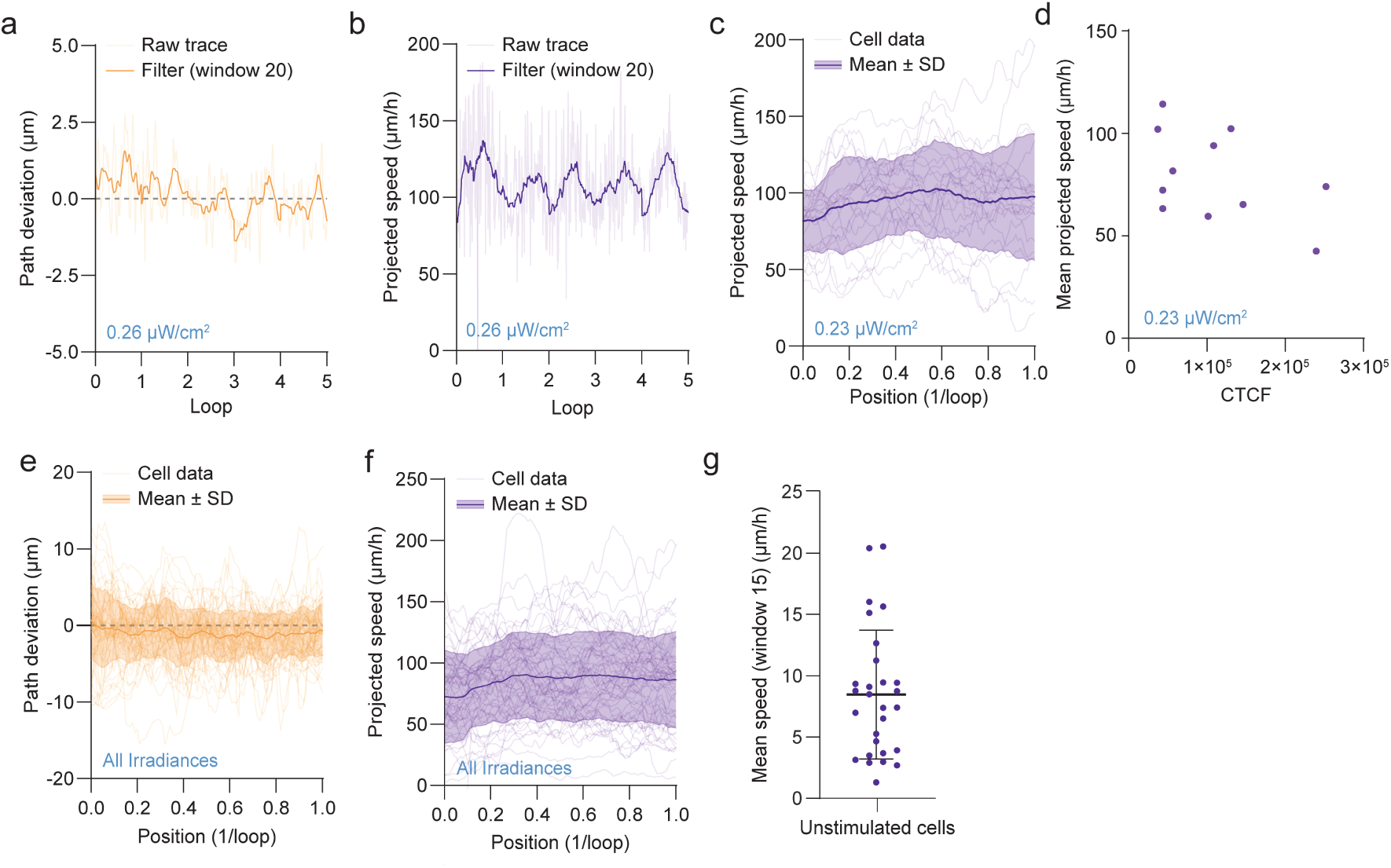
Outcome-driven control of directed cell migration. **a**) The path deviation of a single HT1080-TIAM cell (from Fig. 1c-d) controlled to a circular path for 5 loops, with a constant irradiance of 0.26 µW/cm^2^. **b**) The projected speed of the same cell as in **a**. **c**) The projected speed of HT1080-TIAM cells controlled to circular paths with a constant irradiance of 0.23 µW/cm^2^, separated by individual loops. n = 24 cells. **d**) Average projected speed of cells in **c** over the whole controlled circle path, separated by expression level (corrected total cell fluorescence, CTCF). n = 11 cells. **e**) The path deviation of all HT1080-TIAM cells controlled to circular paths, for all irradiances, separated by individual loops. n = 58 cells. **f**) The projected speed of all cells as in **e**. n = 58 cells. **g**) The mean speed of uncontrolled HT1080-TIAM cells (0 µW/cm^2^). n = 28 cells from 3 separate coverslips.

**Extended Data Fig. 3:**
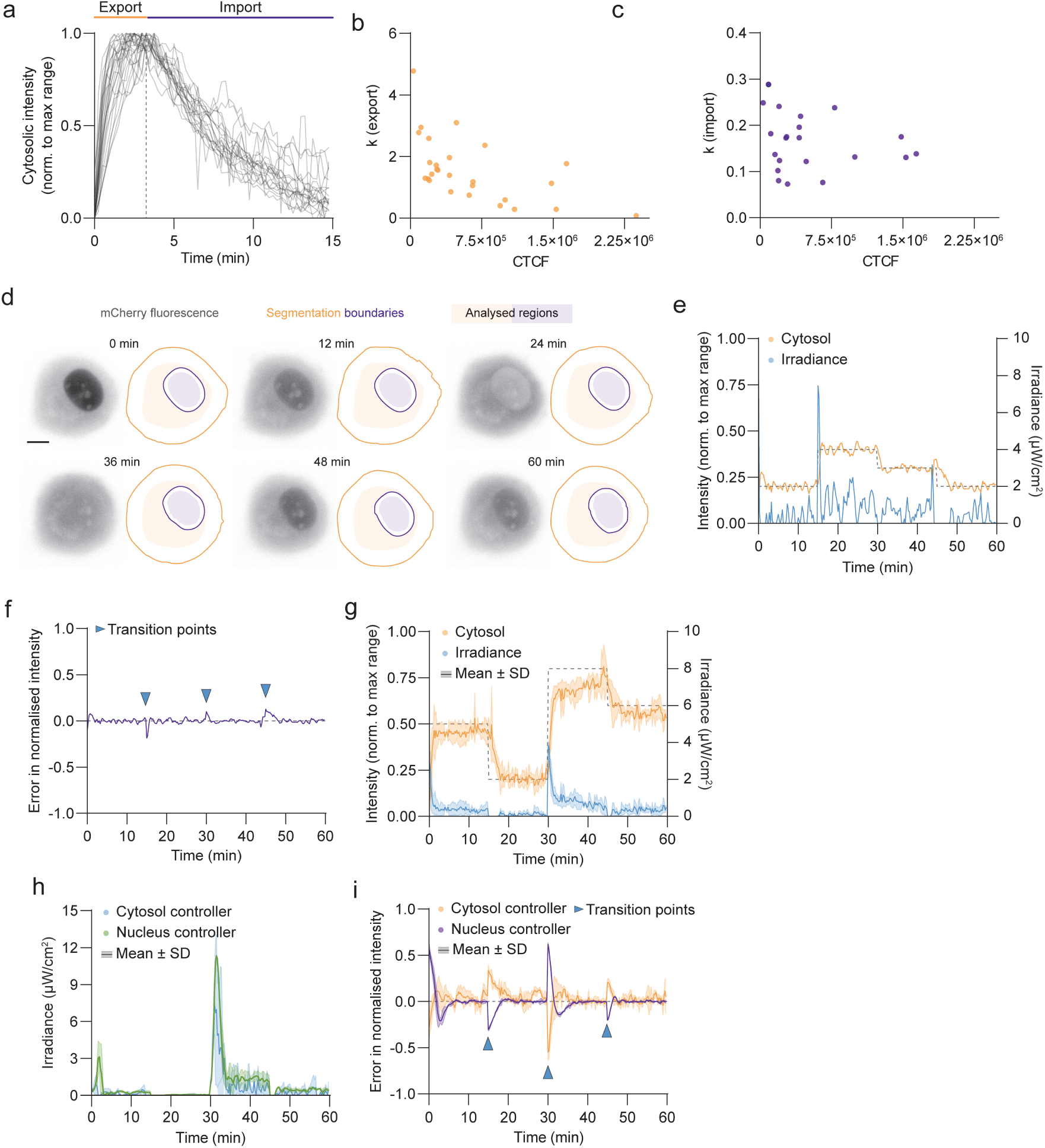
Outcome-driven control of nucleocytoplasmic transport. **a**) Cytosolic intensity of U2OS-LEXY cells illuminated with a fixed irradiance of 10.4 µW/cm^2^ for 3 min (export), and then kept in the dark for 12 min (import). **b**) Decay constant (k) during export for an exponential decay fit to cytosolic intensity during illumination in **a**, separated by expression level (CTCF). **c**) Decay constant (k) during import for an exponential decay fit to cytosolic intensity during the dark state in **a**, separated by CTCF. **d**) A U2OS-LEXY cell (grey) controlled using outcome-driven microscopy to achieve predefined cytosolic intensities, showing segmentation of the cytosol (orange) and the nucleus (purple), and regions analysed for control. Scale bar: 10 µm. **e**) Normalised cytosolic intensity and irradiance of the cell in **d** controlled to various setpoints (dashed line) using a standard PID controller. **f**) Error in normalised intensity for the cell in **d**-**e**, with blue arrowheads to denote changes of setpoint. **g**) Normalised cytosolic intensities and irradiances of multiple cells over time, controlled by a standard PID controller over the whole operating range. n = 4 cells. **h**) Irradiance levels for the cell controlled in Fig. 2f-g, for both the cytosol and nucleus controller. n = 3 runs for each controller. **i**) Error in normalised intensity for the control of cytosol and nucleus intensity in Fig. 2f-g.

## SUPPLEMENTARY INFORMATION

### Supplementary Text

1. Modular Python platform for smart, outcome-driven microscopy
2. Image segmentation for outcome-driven microscopy
3. Controller design for nucleocytoplasmic transport

### Supplementary Video 1

Outcome-driven control of directed cell migration with constant irradiance.

### Supplementary Video 2

Outcome-driven control of directed cell migration with changing irradiance.

### Supplementary Video 3

Outcome-driven control of multiple cells, with an active avoidance system to prevent collisions.

### Supplementary Video 4

Outcome-driven control of nuclear intensity at full operating range, repeated three times in the same cell.

## METHODS

### Cell culture

HT1080 (ATCC) and U2OS Flp-in T-Rex (a gift from Alessandro Sartori) cells were cultured in DMEM high glucose medium (Capricorn DMEM-HPSTA) supplemented with 9% foetal bovine serum (Corning, 35_079_CV) and 1% penicillin/streptomycin (GIBCO, 15140-122) at 37°C with 5% CO_2_. All cell lines were routinely screened (every 8–12 weeks) to ensure they were free from mycoplasma contamination.

The HT1080-TIAM cell line was produced for this study using lentiviral transduction. Lentivirus was produced by transfecting 10 cm dishes of HEK293T cells with 15µg of the transfer plasmid pLenti_TIAM-tagRFP-SSPB-P2A-mVenus-iLID-CAAX (a gift from Mathieu Coppey), together with 10 µg psPAX2 lentivirus packaging plasmid and 5 µg lentivirus envelope plasmid (gifts from Didier Trono, Addgene #12260 and #12259, respectively) using 90 µL 1 mg/mL Polyethylenimine Hydrochloride (Polysciences, 24765). Growth media was harvested 24-and 48-hours post-transfection, filtered with a 0.45 µm syringe filter (Corning, 431220) and virus was precipitated in precipitation solution (5X stock solution: 66.6 mM PEG 6000 (Thermo Scientific Chemicals, A17541.0B), 410 mM NaCl (Thermo Scientific Chemicals, 194090010), in ddH2O, pH 7.2). The viral supernatant was then centrifuged for 30 min at 1500 ×g at 4°C, and the pellet resuspended in PBS (Sigma-Aldrich, D8537) for storage at −80°C. To generate HT1080-TIAM stable line, wild-type cells were seeded to a 24-well plate. 24 hours later, the medium was refreshed with DMEM supplemented with 5 µg/mL polybrene (Sigma-Aldrich TR-1003), and 5 µL viral suspension was added. The medium was refreshed after 24 hours, and cells were first selected in complete medium with 20 µg/mL blasticidin (InvivoGen, 29-11-BL), before sorting for double-positivity with Fluorescence-Activated Cell Sorting (FACS) using a FACSAria™ Fusion Flow Cytometer (BD Biosciences).

The U2OS-LEXY stable line was generated using flp/frt recombinase-mediated cassette exchange. Maternal U2OS Flp-in T-Rex were transfected with the donor plasmid pCDNA5-NLS-mCherry-LEXY and pOG44 (Invitrogen), carrying Flp recombinase. Isogenic cells stably expressing LEXY from a doxycycline-sensitive promoter were selected by treatment with 200 µg/ml hygromycin B (InvivoGen, ant-hg-5).

For cell migration experiments, HT1080-TIAM cells were seeded to 25 mm coverslips (VWR, 631-0172) coated with 200 µL 25 µg/mL fibronectin (Sigma-Aldrich, F1141) at low density, 6-18 hours prior to imaging. 2 mM thymidine (Calbiochem, 6060) was added at the point of seeding to prevent division, and thymidine was kept in the medium for imaging. For nucleocytoplasmic transport experiments, U2OS-LEXY cells were seeded to 25 mm coverslips at medium density, 24 hours prior to imaging. 2 µg/mL doxycycline-hyclate (Abcam, ab141091) was added at the point of seeding to induce expression, and doxycycline was kept in the medium for imaging. Coverslips were mounted in Attofluor cell chambers (Invitrogen, A7816) for imaging.

### Cloning

pCDNA5-NLS-mCherry-LEXY, a mammalian expression vector for LEXY, was derived from NLS-mCherry-LEXY (pDN122, a gift from Barbara Di Ventura & Roland Eils, Addgene #72655), and pCDNA5/FRT/TO (Invitrogen) by conventional molecular cloning.

### Imaging hardware

Imaging was performed using a Nikon Ti dual-turret inverted microscope equipped with a 40× CFI Plan Fluor NA 1.3 oil immersion objective (Nikon, MRH01401), a sample incubator (Tokai-Hit), and a pco.edge cooled sCMOS camera (Excelitas).

For epifluorescence illumination, a pE-4000 LED illumination system (CoolLED) was used as a light source, and ET 514nm Laser Bandpass (Chroma, 49905) and ET mCherry (Chroma, 49008) filter cubes were placed in the lower filter turret.

For optogenetic illumination, a 470 nm LED (Mightex BLS-LCS-0470-15-22) and a Polygon 400 digital mirror device (Mightex) were equipped for the light source, and a 10/90 beamsplitter (Chroma, 21012) was placed in the upper filter turret. Neutral density filters of optical density 0.5 and 0.9 (ThorLabs, NE205B and NE209B) were also placed into the light path to reduce LED intensity. A photodiode power sensor (ThorLabs, S121C sensor with a PM400 console) was used to calculate irradiance at the sample plane. For cell migration experiments, images were acquired with a 60 second interval. For nucleocytoplasmic transport experiments, a 15 second interval was used.

### Smart microscopy platform for outcome-driven microscopy

Custom Python code was written for control of microscope hardware (Supplementary Text 1). We built a modular platform, where an outcome-driven strategy can be defined using separate modules for software-hardware bridging, on-the-fly image analysis, and closed-loop biological control.

In this study, our microscope bridge module uses µ-Manager version 2.0^82^ (Nightly Build 20220930) to control the microscope and peripheral components, and the Pycro-Manager version 0.19.2 package as a translation layer^83^. Our on-the-fly analysis module uses Segment Anything^75^ with custom extensions (Supplementary Text 2), and our closed-loop control module uses either a direct orientation-correction controller (for cell migration), or a proportional-integral-derivative controller (for nucleocytoplasmic transport) (Supplementary Text 3).

### Post-experiment image and data processing

All post-experiment imaging processing was performed using FIJI 1.54f^84^. Data processing was performed using Python, GraphPad Prism 9.5.0, or Microsoft Excel. GraphPad Prism 9.5.0 and Adobe Illustrator 28.7 were used for data visualisation. Matlab v.2023b was used for controller design.

Path deviation was determined by calculating the distance between the centroid of the cell and the nearest point on the path (the projected position) at each timepoint. Projected speed was determined by then dividing the distance between projected points by the imaging interval.

Expression levels of cells were calculated as the corrected total cell fluorescence (CTCF): cell integrated density – (cell area * background mean grey value). Rates of cytosolic intensity change during export or import of LEXY were calculated by measuring cytosolic intensity at 15 s intervals with LED constantly on (10.4 µW/cm^2^) or off, then fitting to a non-linear one phase decay in GraphPad Prism 9.5.0.

For comparison between the standard and gain-scheduled PID controllers, mean error at steady state was calculated by averaging the error from the point where the setpoint was reached. This point is estimated when the derivative of the signal reduced to zero or changed sign. In this way, we clean the error from the transition time.

## CODE AVAILABILITY

Code for our smart microscopy platform, including a guide for integrating outcome-driven microscopy, is available on GitHub: https://github.com/UU-cellbiology/UU_SmartMicroscopy. v.1.0.0 was used for the present study, available at the following link: https://github.com/UU-cellbiology/UU_SmartMicroscopy/releases/tag/v1.0.0.

## DATA AVAILABILITY

The datasets generated and/or analysed during the current study are available on Figshare: https://doi.org/10.6084/m9.figshare.26509702.

## ACKNOWLEDGEMENTS

We thank Eugene Katrukha (Utrecht University) and Dylan Kalisvaart (Delft University of Technology) for helpful discussions, Toni van Capel (Utrecht University) for help with cell sorting, Alessandro Sartori (University of Zurich) for the U2OS Flp-In T-Rex cell line, and Mathieu Coppey, Barbara Di Ventura, Roland Eils, and Didier Trono for sharing plasmids. This work was supported by the Netherlands Organization for Scientific Research (NWO) Gravitation programme IMAGINE! (Project number 24.005.009), by the National Roadmap Initiative NL-BioImaging-AM, and by the Eindhoven, Wageningen Utrecht Alliance through the Centre for Living Technologies.

