## Supplementary Text for "Outcome-Driven Microscopy: Closed-Loop Optogenetic Control of Cell Biology"

#### 1 Modular Python platform for smart microscopy

We developed a Python-based platform to perform automated, outcome-driven experiments. Our platform is organized into multiple modules, as described in Extended Data Figure 1. In this section, we describe the general structure of the platform, followed by a more detailed description of each module. We also describe a small example of how to run an experiment from the user perspective, followed by future improvements and modifications to the platform.

##### 1.1 Modularity and structure

The abstract architecture is as described in Extended Data Figure 1. The user interacts with the platform through a `main.py` script and a graphical user interface (GUI). The `main` connects to the rest of the modules, including the GUI, and runs three parallel, asynchronous loops. The three loops consist of the *interface loop*, the *analysis loop*, and the *control loop*. These loops connect to the independent modules as shown in Fig. 1.

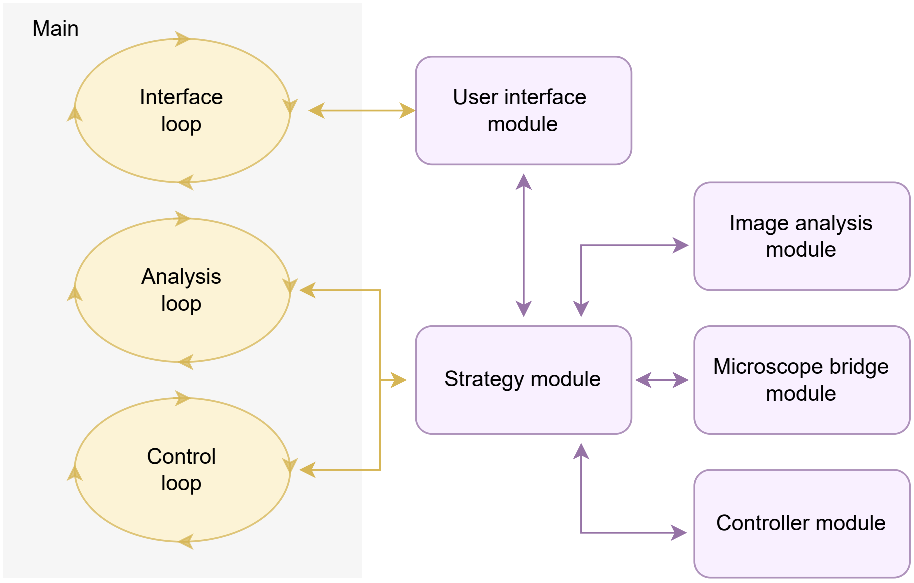


**Fig. 1**: Modules workflow including the `main` and its three internal parallel loops. The modules are the same shown in Extended Data Figure 1.

The modules are imported to the `main` as external libraries based on the user input. Each module is structured as a class with an *abstract class* as a parent, meaning that they must have a list of functions and variables in order to communicate correctly with the rest of the platform. In case the user needs to develop a custom module, it must include these functions and variables as well.

When running an experiment, the platform exports 4 main results. The first result is the time series acquisition, including channels and z-stacks, if previously defined. In addition, the platform exports the cell segmentation for each time step, the controller data (*e.g.*, error, setpoint, laser power) as a `.csv` file, and the metadata of the experiment as a `.json` file.

#### 1.2 Strategy module

The center module of our platform is the (outcome-driven) strategy, *i.e.*, the experiment to be performed. The strategy module can be adapted to any kind of smart microscopy experiment, either outcome-driven, event-driven, or pre-programmed. Here, we show two outcome-driven strategies using optogenetics: one to guide a migratory cell and the other to control the cytosolic and nuclear levels of a cell.

The functions and parameters a strategy module must have, based on their *abstract (template) class* parent, are:

1. `process_step()`: function that connects to the image analysis module and extracts the correct information from the image and segmentation for the closed-loop control.
2. `controller_step()`: function that connects to the controller module and converts the abstract output of the controller to the type of output that will be sent to the microscope.
3. `write_data()`: function to export data to a `.csv` file. The exported data is updated at every step and the variables exported depend on the experiment.
4. `set_parameters()`: function to change internal parameters of the strategy module by the main.
5. `processed_image` and `mod_image`: variables of the segmentation (*i.e.*, output of the image analysis module) and the image to be sent to the modulator, respectively. These variables are retrieved by the main, especially to visualize in the GUI.

For the two outcome-driven strategies presented in the main text, the functions `process_step()` and `controller_step()` take different roles. These functions are called during the analysis and controller loops in `main`, respectively. We describe their specific roles in Table 1.

| | **Cytosol concentration** | **Cell migration** |
|---|---|---|
| `process_step()` | Used to obtain the cell segmentation from a given region. | Used to get directed illumination and for the Active Avoidance of multiple cells. |
| `controller_step()` | Used to obtain the segmentation of the cytosol and nucleus, erode the edges of each mask to avoid overlap, and calculate the concentration of nucleus and cytosol. | Used to update the setpoint if needed and obtain new irradiance from the controller. |

**Table 1**: Roles of the main functions in the strategy module, for the two outcome-driven strategies presented.

In terms of platform, the user can choose any strategy without requirements. The user interface only considers the inputs relevant to the active module, and the other modules are not affected by the strategy used. Furthermore, the strategy module is responsible for and sets the rest of the modules, with the exception of the graphical interface module. If the user wants to develop a new strategy for a tailored experiment, they must build a new strategy module.

However, it is worth noting that our platform does not yet perfectly separate the strategy module from the microscope bridge module explained below. If a smart microscopy strategy requires a completely different experiment design, the microscope bridge must also be adapted. We develop this idea further in Section 1.8.

#### 1.3 Image analysis module

The image analysis module produces a binary mask based on the microscope image. The binary mask represents an area of interest and can be a single cell, a nucleus, a cluster of cells, or sub-cellular structures. For the two outcome-driven examples given, we segment a whole cell for induced cell migration and the cytosol and nucleus of a cell for cytosol concentration.

Although the image analysis module is independent and can be replaced with different segmentation approaches, we used the same algorithm, the Segment Anything Model (SAM), for both examples as the results outperformed any other algorithm we tested. The segmentation is described in detail in Section 2.

The *abstract class* for segmentation defines the following mandatory functions:

1. `setImage()`: function to define the image to segment. This function is typically called after taking an image with the microscope.
2. `setup()`: function to define a region of interest within the image for the segmentation, *e.g.*, where the cell is located. When this function is ignored, the whole image will be used as the region of interest.
3. `run()` and `getResult()`: functions to run the segmentation and to retrieve the result of the segmentation, respectively.

#### 1.4 Controller module

While the strategy module processes the information and decides what variable is optimized, the controller module is where the optimization occurs. The controller module has a single mandatory function, `step()`. On every step in time, this function calculates the signal (*e.g.*, irradiance, area of illumination, frequency) to be sent to the biological sample on the next step. The controller module can be design around any type of controller, from classical linear controllers, to model-based controller, to AI-based controllers. Although the type of controller depends on the experiment, the control module is independent of how the input to the controller (*i.e.*, the signal to be controlled) is obtained. Thus, controller modules are in principle exchangeable.

For the two outcome-driven experiments presented here, we developed two controllers; one for the migration experiment and the other for the cytosol concentration experiment.

##### 1.4.1 Cell migration control

For the cell migration experiment, the `step()` function calculates the *area* of the cell to illuminate with blue light based on the position of the cell and the pre-defined path to follow, always illuminating the edge of the membrane to induce migration in the selected direction (see Figure 1b in the main text). The input then is the raw image of the cell and the (initial) position of the cell.

##### 1.4.2 Cytosol concentration

In turn, for the cytosol concentration experiment, the `step()` function calculates the *irradiance* of the blue light to reach a certain concentration. We use a Proportional-Integral-Derivative (PID) controller to obtain the optimal irradiance value to drive the system to a given setpoint. The input for the controller are the estimated and desired concentration. The controller design is described in detail in Section 3.

#### 1.5 Microscope bridge module

The bridge module is the independent module in charge of communicating with the microscope. Thus, the platform controls the hardware and the time series acquisition through the bridge module. For the two outcome-driven examples given, the bridge module requires access to the camera, the light modulator device, and the blue light LED. Still, for a different experiment, the bridge module may have access to other hardware components such as stage, objectives, or temperature.

For the presented experiments, we controlled a Nikon Ti dual-turret inverted microscope using the Micromanager software [1], a versatile open-software program that allows to control many microscopes from different brands. To communicate with Micromanager, we built a bridge module based on `Pycromanager` [2], a Python-based library that allows easy communication with Micromanager.

Thanks to the modularity design of our platform, the bridge module can be exchanged to run exactly the same experiment in different systems. A bridge module can be developed to use other communication libraries, such as `pymmcore-plus` [3], `navigate` [4], or `ImSwitch` [5]. Furthermore, we are working on developing bridge modules for other (commercial) microscopes. This is described more in depth in Section 1.8.

For any microscope, a bridge module must have the following functions and parameters:

1. `live_image()`: function to take a live image or *snap* from the microscope.
2. `set_modulation()`: function to set a binary mask into the modulator device for the blue-light illumination. This function can be adapted to systems without modulators, *e.g.*, setting a scanning area for a scanning microscope.
3. `run_acquisition()`: function to run the multi-dimensional acquisition. This starts the time-lapse acquisition with the given channels, z-stack, and positions.
4. `shutdown()`: function to start the protocol to turn off the hardware after the experiment, if needed.
5. `Acq_image`: the current image of the multi-dimensional acquisition, once it is running.
6. `modulator_dict`: a dictionary with the information for the modulator device, namely the binary mask and the LED power.
7. `im_height` and `im_width`: are parameters retrieved by main, representing the height and width of the camera images, in pixels.
8. `mod_height` and `mod_width`: are parameters retrieved by main, representing the height and width of the modulator, in pixels.

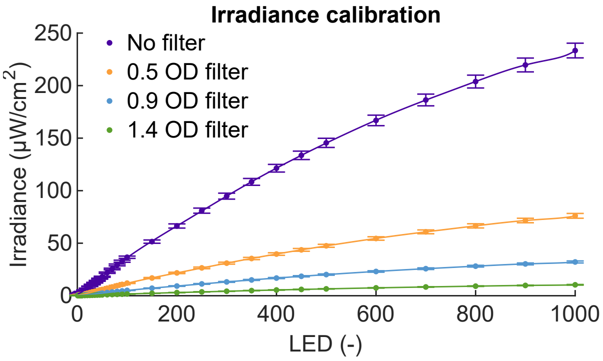


**Fig. 2**: Measured irradiance for different optical filters and a fitted 11-order polynomial.

9. `acq_path` and `acq_name`: are the folder and file names for the resulting time-lapse images. These variables are defined by the user in the main.
10. `image_ready` and `modulator_ready`: are boolean variables observed in main to know if a new image is retrieved or if the modulator image was correctly assigned, respectively.
11. `running`: variable to notify the main that the multi-dimensional acquisition is currently running.

##### 1.5.1 Irradiance calibration

To achieve reproducible control of irradiance-dependent processes, the irradiance at the sample plane was characterized for the Nikon Ti microscope. In this way, the commands generated from the controller are decoupled from the microscope on which the experiment is running. We used a ThorLabs PM400 powermeter with a S121C head placed at the sample plane to measure the irradiance, whilst the commands to the 470 nm illumination LED was varied. We included as well optical density filters to reduce the total irradiance, and calibrate for each filter. The results of this calibration experiment is presented in Figure 2. For each calibration curve, a fitted polynomial was used to convert irradiance targets from the controller to commands for the LED.

#### 1.6 Graphical user interface module

We developed a graphical user interface (GUI) based on the `Tkinter` python library [6]. The user interface is shown in Fig. 3. The GUI has three main functionalities, to **show** the microscope pictures and segmentation, to **interact** with the user via clicks, and to **start** the time series acquisition. The GUI communicates with the segmentation and bridge module to show the pictures and segmentation, obtaining the images from them. When a user interacts with the GUI, the GUI sends the commands to the strategy module. Once the user wants to begin the time series, the *Acquire* button sends the command to the bridge module.

To communicate with the other modules and the `main`, we import an additional python file called `globVars.py` on every module. This file has global variables that are read from multiple classes.

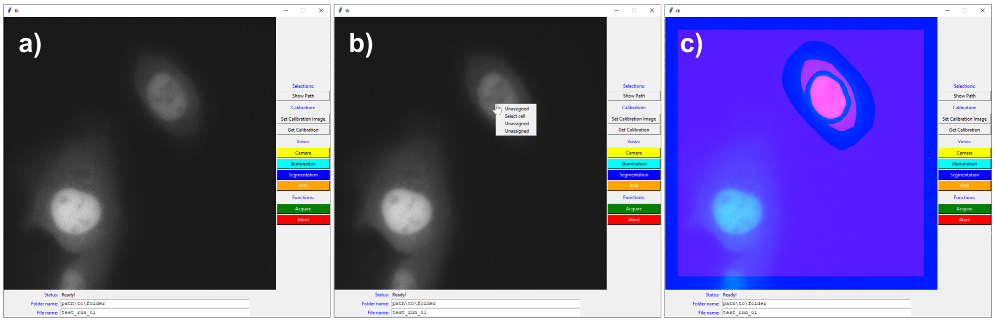


**Fig. 3**: Graphical user interface in three different states. a) when the software is just initialized, b) when right-clicking the cell to segment, including the drop menu, and c) the RGB visualization showing camera image, cell segmentation, and illumination pattern.

#### 1.7 User steps

Here, we give a brief step-by-step for a user to run an outcome-driven experiment using our platform. Let's say the user wants to run the cytosol concentration experiment. After finding and focusing on a single cell, the user needs to go through the following steps:

1. First, the user needs to **define the input variables** in the `inputs.yaml` file. The input variables include the strategy and bridge module to be used, the variable to be controlled by the PID (cytosol or nucleus concentration), the name of the file and folder, and acquisition parameters such as exposure time, acquisition channel, and time series details.
2. With the input variables defined, the user must run the `main`, which opens the GUI. The user then needs to **select the cell** to be controlled, right-clicking the GUI and selecting the segmentation option from the drop menu.
3. Before running the experiment, the user will need to **calibrate the experiment**. The calibration depends on the experiment. For cytosol concentration experiments, we recommend running an initial measurement with a high setpoint to measure the total dynamic range of the cell.
4. Once the user is satisfied with the cell segmentation and calibration, they can start the acquisition by **pressing the `Acquire` button**. In our modules, this will trigger the multi-dimensional acquisition of Micromanager.

#### 1.8 Future modifications

The advantage of modularity is not only that modules are exchangeable, but also that the platform is easily upgraded and improved. Among the improvements planned to the platform, we first intend to **separate further the bridge module with the strategy module**. In particular, the platform is structured around the multi-dimensional acquisition process of Micromanager. This structure is ideal for our outcome-driven experiments and can be translated to other microscopes easily, but it limits experiments to having a constant acquisition configuration, such as time step and number of Z-stack layers. To make our platform flexible, we plan to follow the structure offered by the `pymmcore-plus` library. `pymmcore-plus` is an alternative to `Pycromanager` to control Micromanager-compatible hardware, and is based around a *queue of events* instead of the multi-dimensional acquisition, *i.e.*, a list of orders to give to the microscope.

Furthermore, we plan to **make a new interface** based on the `Napari` library [7]. `Napari` is an extensive visualization library, widely used for microscope data. This new interface offers more functionalities and interaction with the user, and it is also compatible with `pymmcore-plus`.

Additionally, we intend to **develop more bridge modules** to control microscopes that are not compatible with Micromanager. We are especially interested in extending our platform to commercial microscopes such as Zeiss and Leica. For Zeiss microscopes, we plan to use the newly available API functionality, and the LAS X software of Leica also offers opportunities for external control.

Finally, we are aware that many solutions, tools, and architectures are being developed for smart microscopy. For our platform to be compatible with multiple (new and incoming) software and hardware, we intend to **make our bridges portable** with solutions such as `Docker` [8]. `Docker` allows to export *containers*, which are pieces of software that can be used independently of the set of libraries, language, or the computer‘s operative system. Thus, our platform will be compatible with modules created by external community members even if they are not based on Python.

#### 2 Image segmentation

Ass mentioned previously, we used the Segment Anything Model (SAM) for cell segmentation. The Segment Anything project rephrased instance segmentation into a general task by leveraging the masked auto-encoder (MAE) [9] together with an extensive dataset of "over 1 billion masks on 11M licensed and privacy respecting images" [10].

Unsupervised learning strategies, such as MAE's masked-data reconstruction, are shown to generalize well to unseen data [9],[11]. SAM adopted a pre-trained MAE as its image encoder and queries, based on prompts (points or bounding boxes), are decoded by a mask decoder. SAM introduced and trained this mask-decoder, together with components to encode the different types of prompts. A priori, mask-decoding is a relatively simple task, and, especially considering the magnitude of the training dataset, it is unsurprising that this also generalizes well.

This motivates the use of SAM to a wide variety of image types, including phase-contrast or confocal microscopy data. In practice, SAM was sufficiently accurate and robust for our experiments, such that fine-tuning by transfer-learning was unnecessary for our experiments.

##### 2.1 custom interface

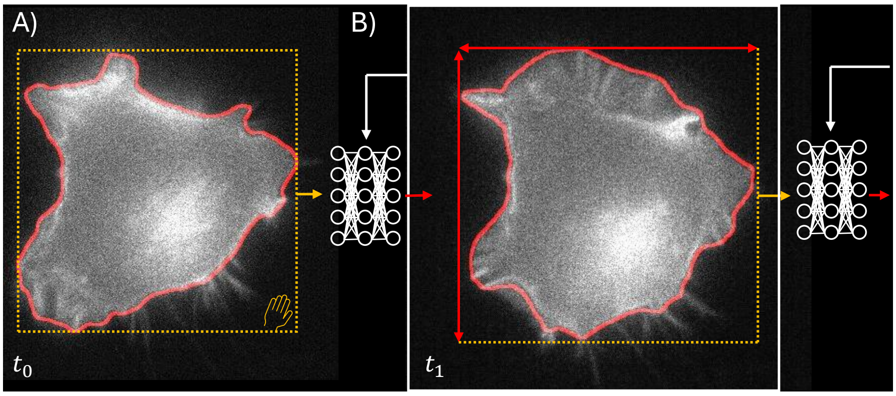


**Fig. 4**: Image segmentation for the migrating cell. A) A bounding box is provided by the user to select a cell and used as a prompt in SAM. A new bounding box is extracted based on the segmentation output. B) The new image is embedded in SAM and prompted using the new bounding box. This process is iterated until the end of the experiment.

SAM and its interface were designed for 1024×1024 8-bit RGB images. Our (8- or 16- bit) single-channel time series are therefore not natively supported. A single frame is therefore re-scaled, repeated 3 times for "RGB" and normalized. The pre-processed image is fed to the model through the (private) tensor functions, bypassing the interface, to preserve the 16-bit accuracy. To initialize the segmentation pipeline, a user specifies the object(s) of interest using a bounding box(es) in the UI (See section 1). Bounding box prompts were chosen as they are inherently more descriptive then a single point (and a single point did not produce robust results during testing). Optionally, the bounding box can be updated to fit the result of the segmentation. The bounding box is re-used for the next image in the sequence and these steps are iterated during the remainder of the experiment (see Fig. 4).

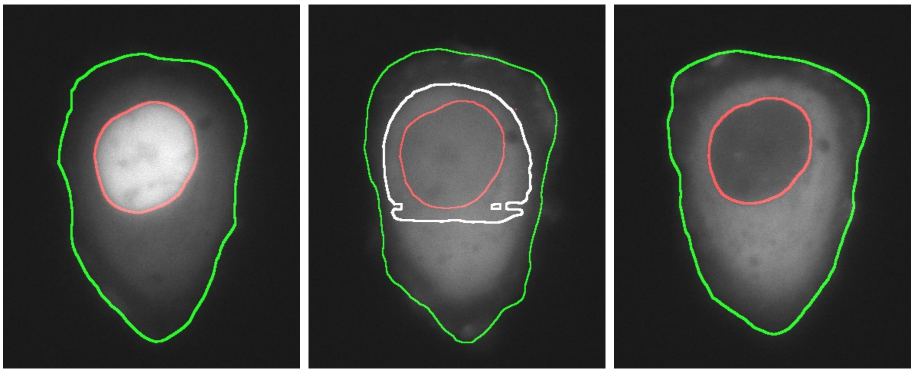

**Fig. 5**: Image segmentation with filtering for LEXY with default, equivalent and inverted Cytosol-Nucleus concentrations. The outline of the cytosol and nucleus segmentations are shown in green and red respectively. For the middle image, the unfiltered nucleus segmentation is shown in white.

During the migrating cell experiments we updated the bounding box using the segmentation outputs. For the nuclear import/export experiments, the bounding box was unchanged during the recordings. This is motivated by the negligible movement of the cells during a recording session, while also preventing undetected problematic timepoints (at which nuclear and cytosol have nearly equivalent intensities) from effecting the bounding box. Problematic timepoints are detected by nucleus size (more than ±20% change w.r.t. previous nucleus), nucleus position (less than 85% overlap with previous nucleus) or morphology (circularity smaller than 0.85). For problematic timepoints, the segmentation results are replaced by results from the previous timepoint to perform image analysis and control (see Fig. 5).

### 3 Controller design for nucleocytoplasmic transport

To actively control the nucleocytoplasmic concentrations, we aim to derive a target illumination intensity from a measured fluorescence intensity, with robustness against cell-cell variations. As mentioned, we employed a PID controller to illustrate the efficacy of simple techniques [12]. Although PID control is rooted in linear system theory, its versatility leads to straightforward adaptations to deal with the nonlinear dynamics in nucleocytoplasmic transport.

We first derive a first-principle model for the measured intensity dynamics, which is then used to find suitable PID parameters. The nucleic and cytosolic intensity measurements $i_\bullet$ are quantified in terms of the saturation point $s$ for ease of user interaction. The saturation is computed from the minimum and maximum achievable concentration $i_{\bullet,\min}$ and $i_{\bullet,\max}$,

$$s = n(i_\bullet) = \frac{i_\bullet - i_{\bullet,\min}}{i_{\bullet,\max} - i_{\bullet,\min}}, \tag{1}$$

were we define $n(i_\bullet)$ as the **saturation map function**. These minimum and maximum intensities are derived experimentally by exposing the cell to a constant irradiance of 10.4 µW cm$^{-2}$ until no further export was observed. This step corresponds to the *calibration* step mentioned on the user guide in Sec. 1.7.

#### 3.1 System modelling and identification

The four different states of the LEXY construct led to a four-compartment model, presented in Fig. 6a. In the figure, $n$ and $c$ are respectively the concentrations in the nucleus and cytosol, with sub-indexes $i$ and $a$ showing (optogenetically) inactive or active proteins, respectively. These four types of concentration represent four states, with specific transition rates $k$ between them. Additionally, the term $u$ represent the irradiance of the optogenetic light, affecting the transfer rate between active and inactive states.

The transition rates $k_d, k_{ex}$ and $k_{im}$ are constant, whilst we assume a linear relationship of illumination intensity $u$ on the activation rates $k'_n$ and $k'_c$, $k'_n = k' + k_n u$ and $k'_c = k' + k_c u$. Experimentally, we found that the nucleic and cytosolic intensities increased uniformly over time, indicating that diffusion dynamics are of a different time-scale compared than the transition rates between states, allowing us to model the dynamics as reaction rate equations.

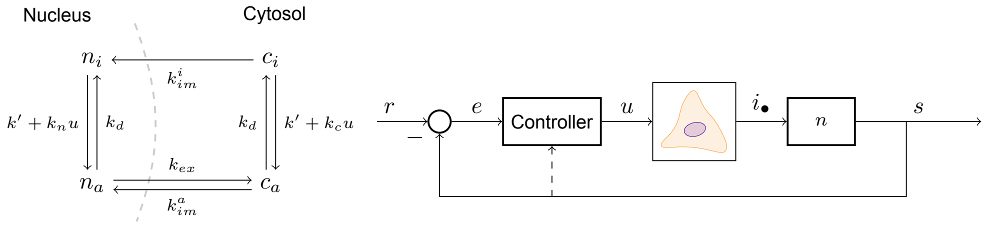


(a) (b)

**Fig. 6**: **(a)** Four-compartment model of light-induced nucleocytoplasmic transport with transition rates. **(b)** Block diagram of signal propagation, how a given target saturation setpoint $r$ is controlled using measured intensities $i_\bullet$ using a PID controller, with the saturation map $n$.

Combining the rate equations with a linear measurement model for the intensity for the nucleocytoplasmic concentrations, leads to

$$\begin{bmatrix}\dot{n}_a\\ \dot{n}_i\\ \dot{c}_a\\ \dot{c}_i\end{bmatrix} = \underbrace{\begin{bmatrix}-k_d-k_{ex} & k' & k^a_{im} & 0\\ k_d & -k' & 0 & k^i_{im}\\ k_{ex} & 0 & -k_d-k^a_{im} & k'\\ 0 & 0 & k_d & -k'-k^i_{im}\end{bmatrix}}_{A}\begin{bmatrix}n_a\\ n_i\\ c_a\\ c_i\end{bmatrix} + u\underbrace{\begin{bmatrix}0 & k_n & 0 & 0\\ 0 & -k_n & 0 & 0\\ 0 & 0 & 0 & k_c\\ 0 & 0 & 0 & -k_c\end{bmatrix}}_{B}\begin{bmatrix}n_a\\ n_p\\ c_a\\ c_p\end{bmatrix},$$

$$\underbrace{\begin{bmatrix}i_n\\ i_c\end{bmatrix}}_{y} = \underbrace{\begin{bmatrix}M_n & M_n & M_{nc} & M_{nc}\\ 0 & 0 & M_c & M_c\end{bmatrix}}_{C}\begin{bmatrix}n_a\\ n_i\\ c_a\\ c_i\end{bmatrix}, \tag{2}$$

with $M_n$. $M_c$, and $M_{nc}$ the proportional relation between concentration and fluorescent intensity.

Writing $y_s$ for measurements at time step $s$, and $x$ for the underlying state, we solve

$$\begin{aligned}\min_{\theta, x_0} \quad & \sum_s \|y_s - \hat{y}_s(\theta, x_0)\|_2\\ \text{s.t.} \quad & \hat{y}_s = f(k, u, \theta, x_0)\end{aligned}$$

using Matlab's nonlinear grey-box system identification tools to find the system parameters $\theta = \{k_d, k_{ex}, \ldots, M_c\}$ and initial condition $x_0$ [13, 14]. The function $f$ is the solution of Equation 2 at time step $s$, given the input sequence $u$, parameters $\theta$ and initial condition $x_0$. For numerical stability of the optimization problem, the intensity measurements are normalized to the nucleus intensity, making the maximum nucleic intensity during optimization $\bar{i} = 1$ and the background intensity corresponding to

$\underline{i} = 0$. Note that this normalization differs from the saturation map in Expression 1. The coordinate transform leaves the matrices $A$, $B$ and $C$ unaffected, and the initialization for the parameters in the nonlinear optimization were found heuristically.

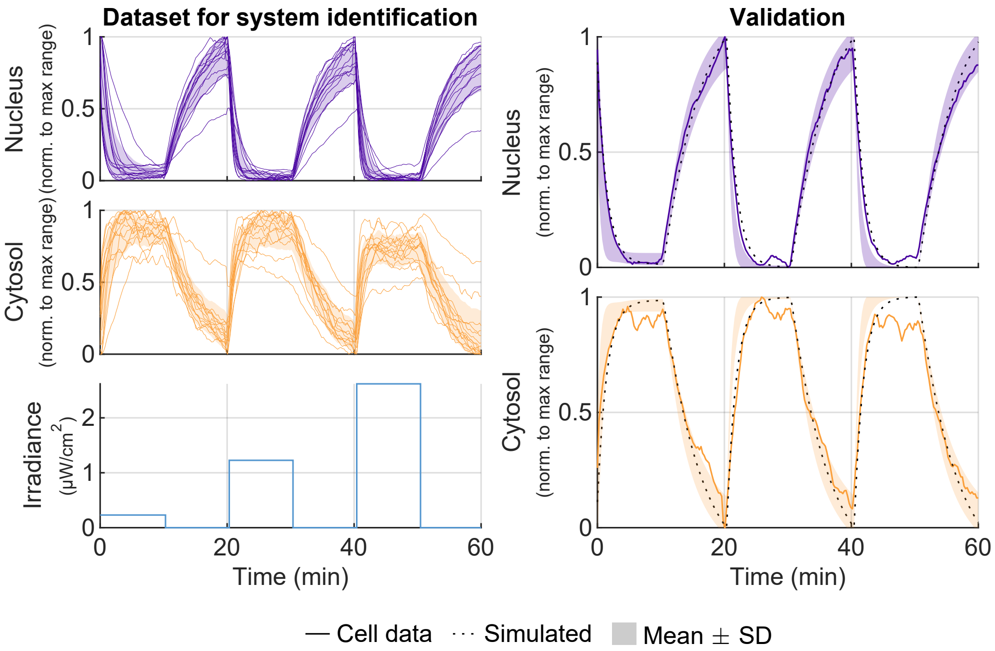


**Fig. 7**: Dataset of 14 sequences used for system identification, for which the standard deviation at each time instance is shaded. For the validation dataset, the standard deviation in dynamics between 8 sampled models is indicated and compared to the model and measurements of a single cell.

For the fitting data, 14 cells were exposed to a 10-minute step sequence of increasing light intensity, $(0.23, 1.23, 2.46)$ $\mu\mathrm{W\,cm^{-2}}$, taking a measurement every 15 seconds. The measured response is illustrated in Figure 7. The optimization problem was then solved for each individual cell, resulting in a range of parameters, of which 8 models with a variance accounted for (VAF) of above 70 % were selected [15]. These models had a variation summarized in Table 2. Together, the selected models had a VAF of $90 - 100$ % and $75 - 95$ % for nucleic and cytosolic intensity respectively. One model, with a VAF of 98 % and 94 % respectively, is presented in Figure 7.

#### 3.2 Controller design

To design the PID controller, the aim was to achieve minimal overshoot ($< 5\%$) across different cells with a response time in the range of 1-5 minutes. With appropriate proportional ($K_p$), integral ($K_i$) and derivative ($K_d$) gains, the controller can achieve reference tracking by driving an error $e_s$ to 0 over time. To steer the intensity $i_\bullet$ to

| | Fit |
|---|---|
| $k_{ex}$ | $0.004 \pm 0.001$ |
| $k_{im}^a$ | $0.2 \pm 0.1$ |
| $k_{im}^i$ | $0.0026 \pm 0.0009$ |
| $k_d$ | $0.0011 \pm 0.0003$ |
| $k'$ | $0.04 \pm 0.02$ |

| | Fit |
|---|---|
| $k_n$ | $5 \pm 4$ |
| $k_c$ | $0.019 \pm 0.007$ |
| $M_n$ | $0.03 \pm 0.01$ |
| $M_c$ | $0.16 \pm 0.04$ |

**Table 2**: Mean fitted parameters for the nucleocytoplasmic transport dynamics with their standard deviation.

| Nucleic saturation $s$ | $K_p$ | $K_i$ | $K_d$ |
|---|---|---|---|
| 0.0 | 90 | 0.5 | 3 |
| 0.25 | 45 | 0.25 | 1.5 |
| 0.5 | 9 | 0.05 | 0.3 |
| 0.75 | 5 | 0.05 | 0.2 |
| 1.0 | 3 | 0.02 | 0.2 |

**Table 3**: Gain-scheduling PID parameters across the operating range for nucleic saturation.

a reference $r_s$, we have $e_s = (i_\bullet)_s - r_s$. In discrete time, the PID controller computes the light intensity $u_s$,

$$u_s = K_p e_s + K_i \sum_{i \leq k} e_i \ \mathrm{d}t + K_d \frac{e_s - e_{s-1}}{\mathrm{d}t}.$$

The integrator component, $\sum e_i \ \mathrm{d}t$, was bounded from below to $-0.1$ to mitigate windup due to a bounded input. To find the controller gains satisfying the requirements, the dynamics was linearized around saturation operating points $n(i_\bullet) \in \{0, 0.25, 0.5, 0.75, 1.0\}$. Integral gains were then maximized until the 5% overshoot limit was reached for the standard deviation in the models in frequency-domain. Next, the proportional gains were increased to increase the response time until the same 5% overshoot limit was reached. Stricter overshoot requirements can be obtained using the same method, at the cost of a slower response time.

For the non-gain-scheduled controller, the controller at saturation $n(i_\bullet) = 0.5$ was chosen, whose gains were later adapted slightly to reduce oscillations at low-export conditions. This lead to the parameter set $(K_p, K_i, K_d) = (9.0, 0.005, 0.3)$

Because of the nonlinear dynamics, gain-scheduling was employed to improve the performance at low and high saturation. Different parameters were found at 4 additional operating points, leading mostly to improved performance at low-saturation conditions. These gains are presented in Table 3.

The linearized closed-loop frequency response from reference input to measured output at different operating points is presented in Figure 8. Note that the PID does not achieve the 5% overshoot requirement when constraint to the response-time wishes across the saturation range, contrary to gain-scheduling.

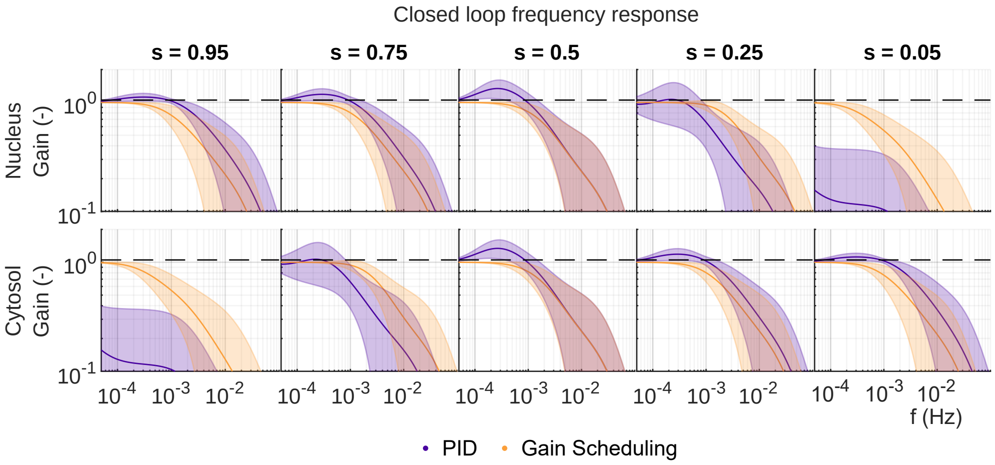


**Fig. 8**: Mean frequency response from reference setpoint to measured output for linearizations of the identified models, the standard deviation between cell models is indicated by the shaded regions. The dashed line indicates the maximum 5 % overshoot limit.

[6] Tkinter - python interface to tcl/tk. URL https://docs.python.org/3/library/tkinter.html.

[7] Sofroniew, N. *et al.* napari: a multi-dimensional image viewer for Python. URL https://github.com/napari/napari.

[8] Overview of docker build (2024). URL https://docs.docker.com/build/.

[9] He, K. *et al.* Masked Autoencoders Are Scalable Vision Learners (2022). URL https://ieeexplore.ieee.org/document/9879206/.

[10] Kirillov, A. *et al.* Segment Anything (2023). URL https://ieeexplore.ieee.org/document/10378323/.

[11] Radford, A. *et al.* Learning transferable visual models from natural language supervision (2021). URL https://proceedings.mlr.press/v139/radford21a.html.

[12] Ogata, K. *Modern control engineering* Instrumentation and controls series (Prentice Hall, 2010).

[13] Ljung, L. *System identification: Theory for the user* Prentice Hall information and system sciences series (Prentice Hall PTR, 1999).

[14] The MathWorks Inc. System Identification Toolbox: 23.2 (r2023b) (2024). URL https://www.mathworks.com.

[15] Verhaegen, M. & Verdult, V. *Filtering and System Identification: A Least Squares Approach* (Cambridge University Press, 2007).
